# A machine learning framework for extracting information from biological pathway images in the literature

**DOI:** 10.1101/2024.06.01.596964

**Authors:** Mun Su Kwon, Junkyu Lee, Hyun Uk Kim

**Affiliations:** Department of Chemical and Biomolecular Engineering, Korea Advanced Institute of Science and Technology (KAIST), Daejeon 34141, Republic of Korea; Graduate School of Engineering Biology, KAIST, Daejeon 34141, Republic of Korea; BioProcess Engineering Research Center and BioInformatics Research Center, KAIST, Daejeon 34141, Republic of Korea

**Keywords:** literature mining, information extraction, metabolic engineering, biological pathway images, object detection

## Abstract

There have been significant advances in literature mining, allowing for the extraction of target information from the literature. However, biological literature often includes biological pathway images that are difficult to extract in an easily editable format. To address this challenge, this study aims to develop a machine learning framework called the “Extraction of Biological Pathway Information” (EBPI). The framework automates the search for relevant publications, extracts biological pathway information from images within the literature, including genes, enzymes, and metabolites, and generates the output in a tabular format. For this, this framework determines the direction of biochemical reactions, and detects and classifies texts within biological pathway images. Performance of EBPI was evaluated by comparing the extracted pathway information with manually curated pathway maps. EBPI will be useful for extracting biological pathway information from the literature in a high-throughput manner, and can be used for pathway studies, including metabolic engineering.

## 1. Introduction

Among the literature across diverse disciplines, a key characteristic of the literature on biology and biotechnology is the visual presentation of complex biological pathways through images (i.e., figures) (Pavlopoulos et al., 2015; Villaveces et al., 2015). Such visual presentations correspond to the key summary of main findings in their respective studies. Thus, curation of biological pathways is crucial for understanding molecular processes in biology and biotechnology such as metabolic engineering (Keller et al., 2022; Sharma et al., 2024) and biomedical sciences (Kuenzi and Ideker, 2020; Lee et al., 2023). In metabolic engineering, for example, metabolic pathways provide information about genes, enzymes, metabolites, and/or other factors involved across the biochemical reactions, which serve as a blueprint for identifying genetic manipulation targets to enhance chemical production. In this context, metabolic databases such as MetaCyc (Caspi et al., 2020), WikiPathways (Pico et al., 2008), Reactome (Fabregat et al., 2017) and KEGG (Kanehisa and Goto, 2000) are useful resources, offering extensive details on biological pathways. These databases continue to be updated, some of them through manual curation. Here, availability of automated extraction of biological pathway information can be useful for pathway studies, offering information complementary to that from the relevant metabolic databases.

There have been remarkable advances in literature mining to retrieve target information from the literature (Gonzalez et al., 2016; Zhao et al., 2021). However, to the best of our knowledge, despite advances in data mining and the importance of biological pathway images in the literature, extracting pathway information from images has not been sufficiently conducted. Extracting target information from images is considered more challenging than text mining due to more variations in visual presentations of biological pathways across different papers. Being able to efficiently extract biological pathway information from a large volume of literature in a standardized, editable format would allow many applications: for example, visual summarization of the up-to-date findings and design of microbial biosynthetic pathways for a target chemical. Object detection models could serve as a potential solution that can recognize and extract target objects (e.g., arrows and texts) within digital images. Examples include DEtection TRansformer (DETR) (Carion et al., 2020) and Faster R-CNN (Ren et al., 2016), both of which detect specific target objects (e.g., arrows in a biological pathway image) in an image. OChemR (https://github.com/markmartorilopez/OChemR) is a comprehensive framework specifically designed to process images of chemical reactions by using DETR. A seeming limitation of OChemR is its unability to handle reaction reversibility (i.e., recognition of only unidirectional arrows, not bidirectional ones) and its ability to detect only molecular reactions presented in a very specific format. Another technology available for extracting information from images is optical character recognition (OCR), which detects texts within images, and transforms them into editable and searchable texts (Hom et al., 2022; Soeno et al., 2024).

In this study, we developed a machine learning framework for extracting biological pathway information (EBPI) from the literature. This framework processes biological pathway images by detecting arrows and texts (e.g., genes, proteins and/or metabolites), and generates biochemical reactions in an editable, tabular format. Here, EBPI employs third-party machine learning models, Faster R-CNN (Ren et al., 2016) and PaddleOCR (https://github.com/PaddlePaddle/PaddleOCR) to detect arrows and texts within pathway images, respectively. To develop and validate EBPI, datasets were prepared from multiple sources and via extensive dataset preprocessing (Fig. 1). Extraction performance of EBPI was examined in comparison with images extracted from 74,853 papers searched for 466 chemicals. Also, EBPI demonstrates its potential with uncovering biochemical reactions not cataloged in well-known biological pathway databases such as MetaNetX (Caspi et al., 2020) and PubChem (Kim et al., 2016). As a literature mining tool, EBPI will contribute to providing important insights into biological pathways of interest based on pathway images extracted from a large volume of literature.

**Fig. 1.**
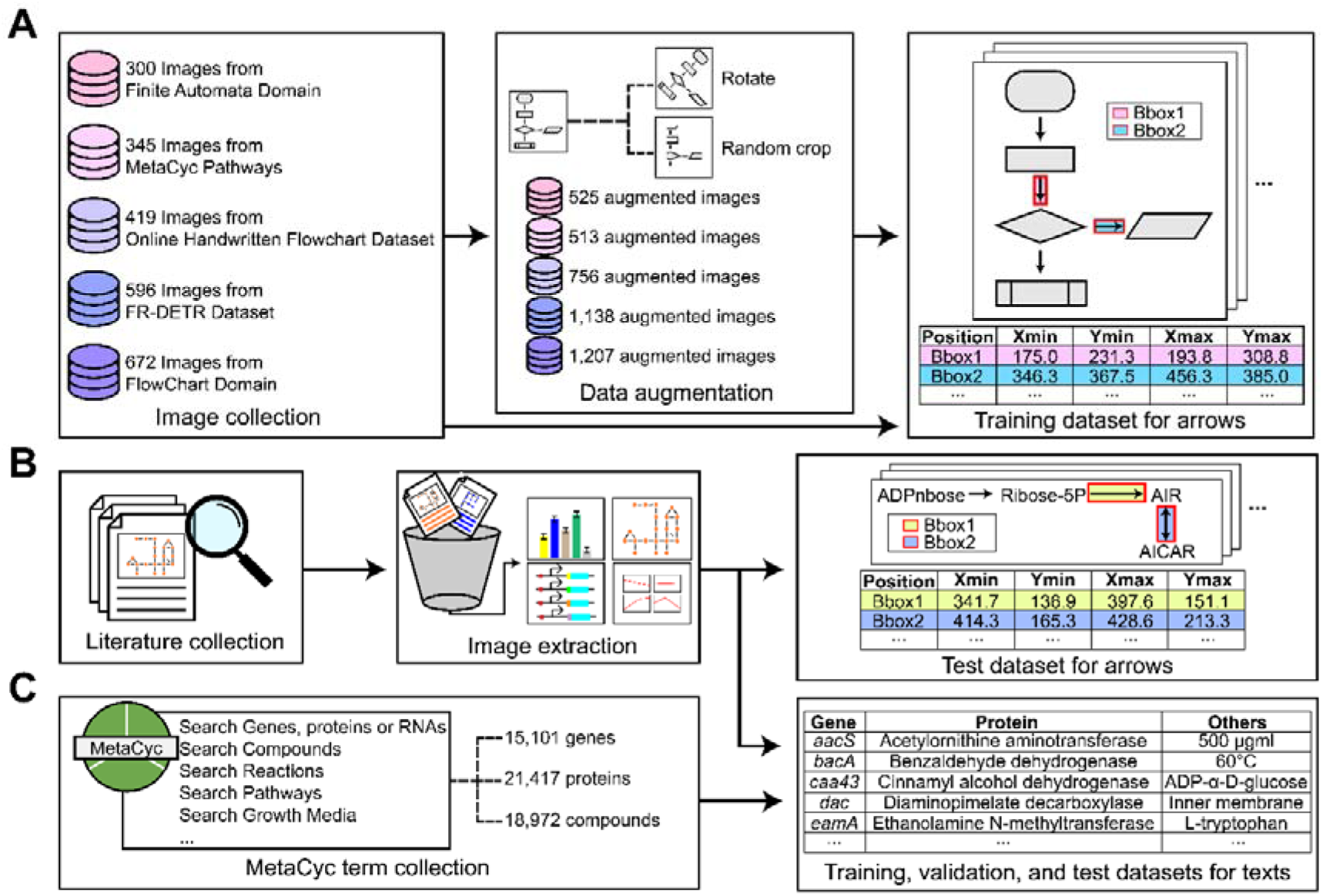
Dataset preparation to develop the machine learning framework EBPI. (A) Preparation of a training and validation dataset for detecting arrows in images. A total of 2,332 images were initially collected from Finite Automata Domain (Bresler et al., 2016), MetaCyc pathways (Caspi et al., 2020), Online Handwritten Flowchart Dataset (https://tc11.cvc.uab.es/datasets/OHFCD_1), FR-DETR Dataset (Sun et al., 2022), and FlowChart Domain (Bresler et al., 2016). Next, 4,139 images were additionally generated from the 2,332 images using rotation and random cropping for data augmentation. The resulting 6,471 images and 43,552 bounding boxes (e.g., “Bbox1” and “Bbox2” in the figure) were used as inputs to detect arrows across the images. (B) Preparation of datasets for detecting arrows and texts within images from papers searched through PMC. Here, 100 biological pathway images were extracted from 89 papers. (C) Preparation of datasets for detecting texts within pathway images from MetaCyc. A total of 59,370 MetaCyc terms, including names of genes, proteins, and metabolites, were collected to prepare training, validation, and test datasets for text detection. These MetaCyc terms were categorized as “gene”, “protein” or “others”.

## 2. Results

### 2.1. Dataset preparation for detecting arrows and classifying texts

Faster R-CNN (Ren et al., 2016) and BioBERT (Lee et al., 2020) within EBPI were trained to detect arrows and classify the detected texts, respectively, within biological pathway images. PaddleOCR, an optical character recognition model, was used to detect texts before implementing BioBERT, but it does not require training. EBPI first identifies biochemical reactions based on the positions of arrows and texts. EBPI uses Faster R-CNN, an object detection model, for bounding box regression of arrows on given images; bounding boxes are a rectangular frame used to define the location of a target object within an image. To train Faster R-CNN model, we first collected a total of 2,332 unique images from five different sources (Fig. 1A and https://zenodo.org/records/11075692): 300 images from Finite Automata Domain (Bresler et al., 2014); 345 images from MetaCyc (Caspi et al., 2020); 419 images from Online Handwritten Flowchart Dataset (https://tc11.cvc.uab.es/datasets/OHFCD_1); 596 images from FR-DETR Dataset (Sun et al., 2022); and 672 images from FlowChart Domain (Bresler et al., 2016). These collected images contain biological and non-biological diagrams with different forms of computer-generated or hand-written arrows. We additionally generated 4,139 images through data augmentation by rotating or cropping the original images. Preparation of the original and the augmented images led to a total 6,471 images. These images are labeled data, having bounding boxes of 43,552 arrows, and they were used as training and validation datasets.

Furthermore, we additionally obtained 100 pathway images through paper searches at PubMed Central (PMC) (Roberts, 2001) using the search term “Metabolic Engineering”, and manually labeled the images to create an external test dataset (Fig. 1B, and https://zenodo.org/records/11075692). These 100 images from PMC appeared to have 1,139 arrows, and they were used to test the model.

Texts detected by PaddleOCR within EBPI were classified into one of three categories (i.e., “gene”, “protein” and “others”) using BioBERT, a pre-trained natural language model designed for biomedical literature. To fine-tune BioBERT, we first collected MetaCyc terms (Caspi et al., 2020) that correspond to genes, proteins, or compounds (Fig. 1C). As a result, following text datasets were obtained: 1) 15,101 genes; 2) 21,417 proteins, corresponding to polypeptides or protein complexes; and 3) 18,972 compounds. Also, 623 papers were additionally collected from PMC using the search term “Metabolic Engineering” to augment “others” terms. This step was necessary because “others” terms should cover a range of items, for example “60°C” and “500 μgml”, in addition to compound names from MetaCyc; meanwhile, the “gene” and “protein” terms are relatively well-defined in MetaCyc terms. For this, PaddleOCR was used to detect texts in images from the 623 papers. The detected texts were manually classified into “others” terms. As a result, 3,880 additional terms were added to “others” based on 2,662 images in the 623 papers. Thus, we now have 22,852 terms of “others” by combining the data from MetaCyc and those from the PaddleOCR results. The combined text dataset was split into training (80%), validation (10%) and test (10%) datasets for the model development.

### 2.2. Development of EBPI

EBPI consists of two modules, one for arrow detection and the other one for text detection (Fig. 2). First, the prepared 6,471 images were processed with Faster R-CNN to detect arrow positions and directions. Next, texts located near the head, tail and middle of the detected arrows were retrieved. These texts correspond to substrates and metabolites of a biochemical reaction, as well as its associated gene or protein. These two modules are further elaborated below.

**Fig. 2.**
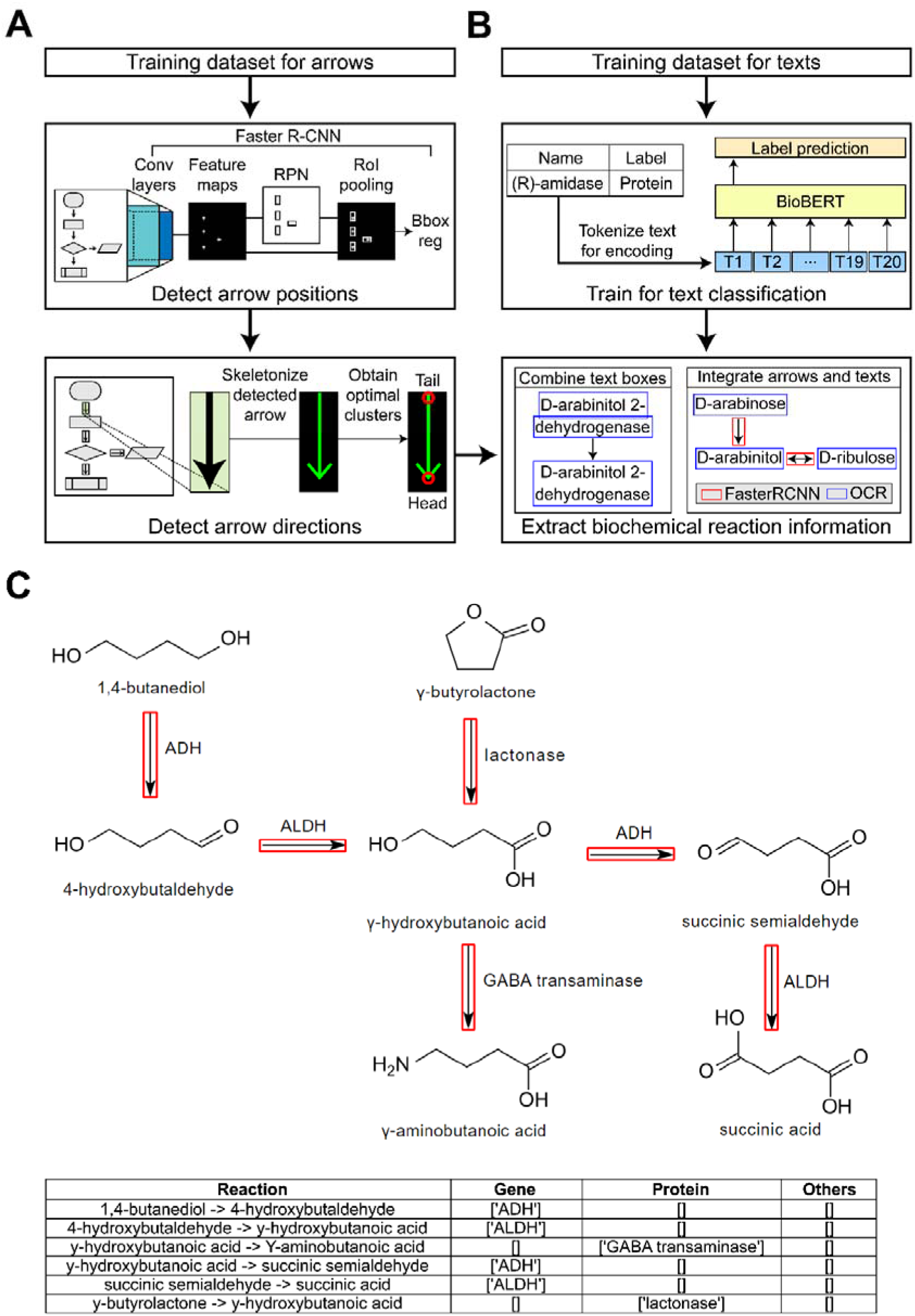
Overview of EBPI development for extracting information on arrows and texts from pathway images. (A) Detection of the position and direction of arrows in the prepared images. Each image was used as input to train Faster R-CNN that detects arrow positions. This process involves bounding box regression (“Bbox reg” in the figure), and utilizes convolutional (Conv) layers, feature maps, a region proposal network (RPN), and region of interest (RoI) pooling (middle box). The detected arrows were subsequently skeletonized, and their end-points were extracted by using OpenCV (lower box). Finally, the end-points were subjected to *k*-means clustering to determine the head and tail of an arrow, thus determining its direction. (B) Detection and processing of text data from images. Each of the 59,370 terms was tokenized into a maximum of 20 tokens. These input tokens were used to train BioBERT, which consequently categorizes the texts into “gene”, “protein” or “others” (middle box). Once images are ready, arrows and texts were detected using Faster R-CNN (“FasterRCNN”) and PaddleOCR (“OCR”) (lower box). Finally, bounding boxes of texts were combined if they overlapped or were in close proximity. This process of the dataset preprocessing was also implemented to prepare the test dataset. (C) An example of using EBPI to extract information from an image that illustrates the metabolic pathway of γ-hydroxybutyric acid (or γ-hydroxybutanoic acid in the image; adopted from https://en.m.wikipedia.org/wiki/File:GHB_metabolic_pathway.svg; December, 2023) by using EBPI.

EBPI uses Faster R-CNN to detect bounding boxes of arrows (Fig. 2A). In order to optimize Faster R-CNN, we tested 32 different hyperparameter sets by varying `batch_size`, `lr`, max_norm, and weight_decay via grid search (Methods). The best-performing model was determined according to its maximum mean average precision (mAP) score. Using the test dataset (i.e., the 100 collected images mentioned above), the best Faster R-CNN model showed the mAP of 0.433, which is competitive when compared to OChemR. After the detection of arrows, their directions were detected next. For this, the detected arrow was first transformed into a morphological skeleton to get end-points (i.e., coordinates of the corners of a bounding box) using thinning, which reveals overall structure of an arrow (i.e., a vertical green line in the third box of Fig. 2A). Finally, the end-points were subjected to clustering to determine the head and tail of the arrow.

Next, we focused on text classification using BioBERT for the detected texts by PaddleOCR (Fig. 2B). We fine-tuned BioBERT to classify the texts into “gene”, “protein” or “others”. In this case, the best-performing model was selected based on the maximum accuracy by changing `batch_size` and `lr` with 12 different hyperparameter sets (Methods). We found that the best BioBERT model showed accuracy of 0.975 using the test dataset. There are two reasons for this very high accuracy. First, most proteins have standardized terms in their names. For example, 18,780 terms contained a suffix “-ase” to indicate enzymes, which correspond to 87.7% of the entire 22,852 protein terms collected. Similarly, 14.3% of protein terms included another protein-indicating term “subunit”. Second, gene names have much shorter character lengths compared to those of proteins and compounds. In the text dataset, the average character length of gene names was 4.517 in comparison with 34.731 and 24.490 for protein names and “others” terms, respectively. The short length of gene names, in addition to their standardized names, might have contributed to a high classification accuracy.

The last step of EBPI is to integrate the captured information from arrows and texts and generate the biochemical reaction information in a tabular format (Fig. 2B,C). During this process, the following were additionally considered.

1. In some cases, the detected texts for a single name were split into multiple lines due to a limited space within a bounding box. Bounding boxes for such texts can overlap slightly in their areas. The intersection of bounding boxes was used to combine separately detected texts into a complete biological name (e.g., metabolite name or gene name) (third box in Fig. 2B).
2. Common chemical formulas such as “OH” and “COOH”’ in molecular structures were excluded during this integration step to prevent these functional groups from being combined with biological names.
3. Texts associated with an arrow were identified by calculating the shortest distances between the arrow’s head/tail and the nearby texts. Texts near the head or tail were not subjected to classification using BioBERT. Text classification was only conducted if a text is located near the middle of an arrow, instead of near its head or tail.

Taken together, EBPI provides information on biochemical reactions from pathway images by automatically integrating detected arrows and texts through its two corresponding modules (Fig. 2C).

### 2.3. Performance of different deep learning models within EBPI for arrow detection

We investigated the classification performance of several object detection models designed for arrow detection (i.e., bounding box regression of arrows) to justify the selection of Faster R-CNN (more precisely, Faster R-CNN V2 with FPN) in this study (Table 1). We used the first and second versions of Faster R-CNN with Feature Pyramid Networks (Faster R-CNN V1 with FPN and Faster R-CNN V2 with FPN, respectively), DETR, and Deformable DETR. In order to quantify the regression performances, the models were trained using the same training and validation datasets via grid search as mentioned above (https://zenodo.org/records/11075692), but each of these models was optimized by using independent hyperparameter sets. Subsequently, using the test dataset, Faster R-CNN V2 showed the best performance in terms of the average precision scores, followed by Faster R-CNN V1, Deformable DETR and DETR (Table 1). We also compared the optimized Faster R-CNN V2 with OChemR for arrow detection using the test dataset (Table 1). The optimized Faster R-CNN V2 again showed the higher average precision scores. This process validates the use of Faster R-CNN within EBPI to detect arrows in pathway images from literature.

**Table 1.** Arrow detection performance of EBPI with different deep learning models.

| Object detection model | mAP | AP<sub>50</sub> | AP<sub>75</sub> | AP<sub>S</sub> | AP<sub>M</sub> | AP<sub>L</sub> |
| --- | --- | --- | --- | --- | --- | --- |
| Faster R-CNN V1 with FPN | 0.396 | 0.760 | 0.364 | 0.327 | 0.400 | 0.421 |
| Faster R-CNN V2 with FPN | **0.433** | **0.778** | **0.429** | **0.330** | **0.432** | **0.612** |
| DETR | 0.310 | 0.626 | 0.269 | 0.170 | 0.328 | 0.496 |
| Deformable DETR | 0.339 | 0.583 | 0.369 | 0.206 | 0.359 | 0.524 |
| OChemR | 0.021 | 0.066 | 0.006 | 0.006 | 0.032 | 0.023 |

The best performance is presented in bold. mAP indicates the mean average precision calculating the average precision at different intersection over union (IoU) values, ranging from 0.50 to 0.95. AP_50_ and AP_75_ correspond to the average precision scores at IoU values of 0.50 and 0.75, respectively. AP_S_, AP_M_ and AP_L_ represent the average precision scores for objects of small (under 32 × 32 pixels), medium (between 32 × 32 and 96 × 96 pixels) and large (over 96 × 96 pixels) sizes, respectively. Also, OChemR has object detection functionality, but it is not an object detection model itself.

### 2.4. Extraction of biochemical reaction information for 466 target chemicals

After its development, EBPI was tested to evaluated its performance in extracting biochemical reaction information for a range of target chemicals producible through metabolic engineering (Figs. 3 and 4). A comprehensive bio-based chemicals map was recently released, which presents biological and/or chemical pathways for the production of 466 target chemicals (Jang et al., 2023). Thus, we decided to extract biochemical reaction information from a large volume of the literature for these 466 chemicals, and to compare this retrieved information with the reaction data presented in the map (Fig. 3A). This comparison would allow evaluating and validating the extraction performance of EBPI for industrially important chemicals.

**Fig. 3.**
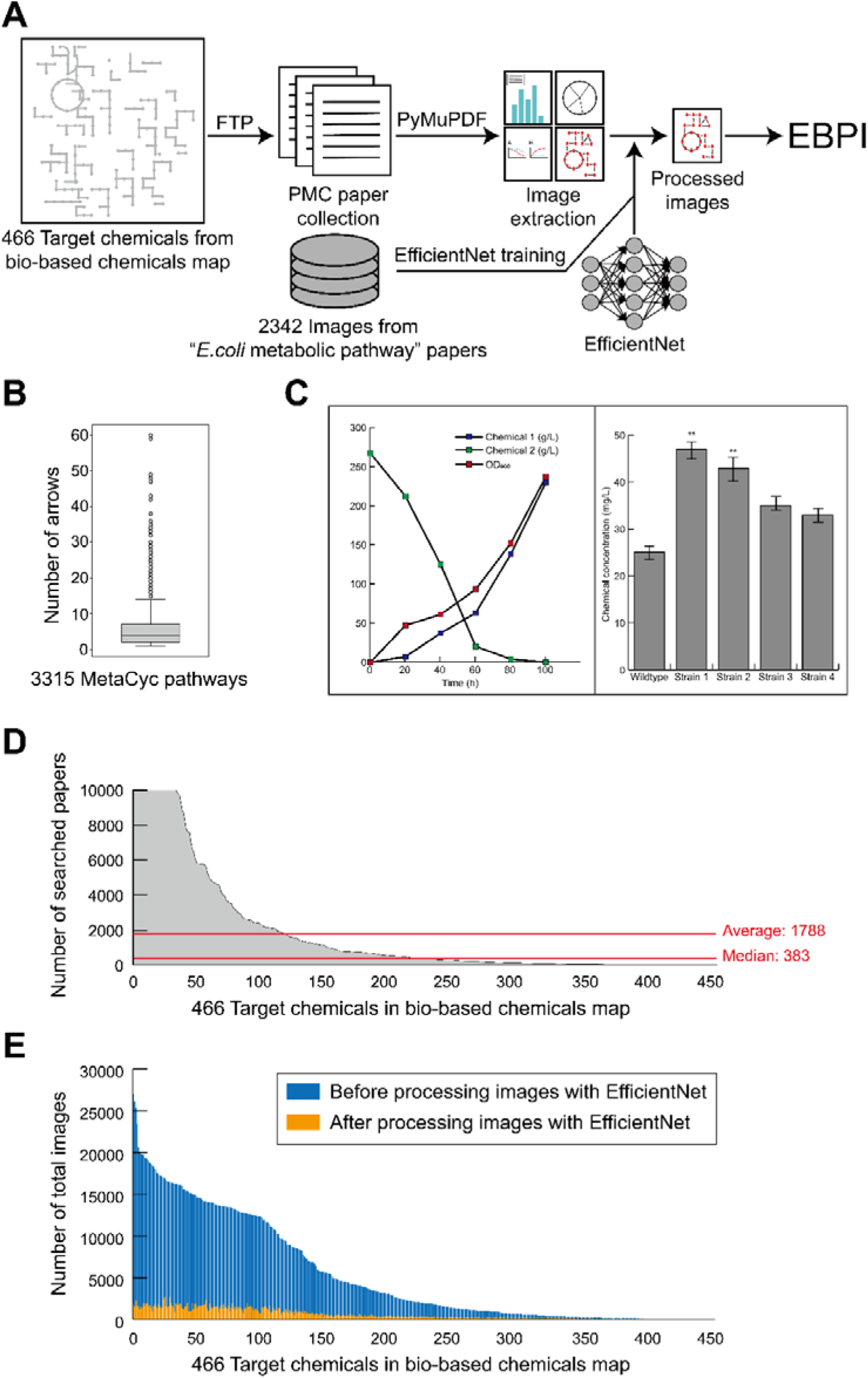
Preparation of images from the literature illustrating biological pathways for 466 target chemicals derived from the bio-based chemicals map (Jang et al., 2023). (A) Preparation of biological pathway images using PyMuPDF and EfficientNet for EBPI implementation. Papers were collected from the NCBI FTP site for the 466 target chemicals, and pathway images were extracted from the collected papers by using PyMuPDF. EfficientNet was next used to select true “pathway” images, while discarding “non-pathway” images. (B) Box plot indicating the distribution of arrow counts retrieved from 3,315 MetaCyc pathway maps. With an average of 5.76 and a median of 4.00 arrows per pathway image, pathway images having five or greater arrows were considered as “pathway” images (positive data), while those having fewer than five arrows (negative data) were mostly considered as “non-pathway” images. The negative image data further underwent manual curation. Both positive and negative image data were used to train EfficientNet. (C) Examples of non-pathway images that were initially extracted from the papers. (D) Number of papers searched for each of the 466 chemicals from the bio-based chemicals map. A maximum threshold of 2,000 papers per chemical was set to retrieve pathway images. (E) Numbers of pathway images before and after treatment with the trained EfficientNet. Only the images (in orange) that passed the EfficientNet-based selection were considered as inputs for EBPI. In this study, 49,846 images were finally selected for 421 chemicals.

**Fig. 4.**
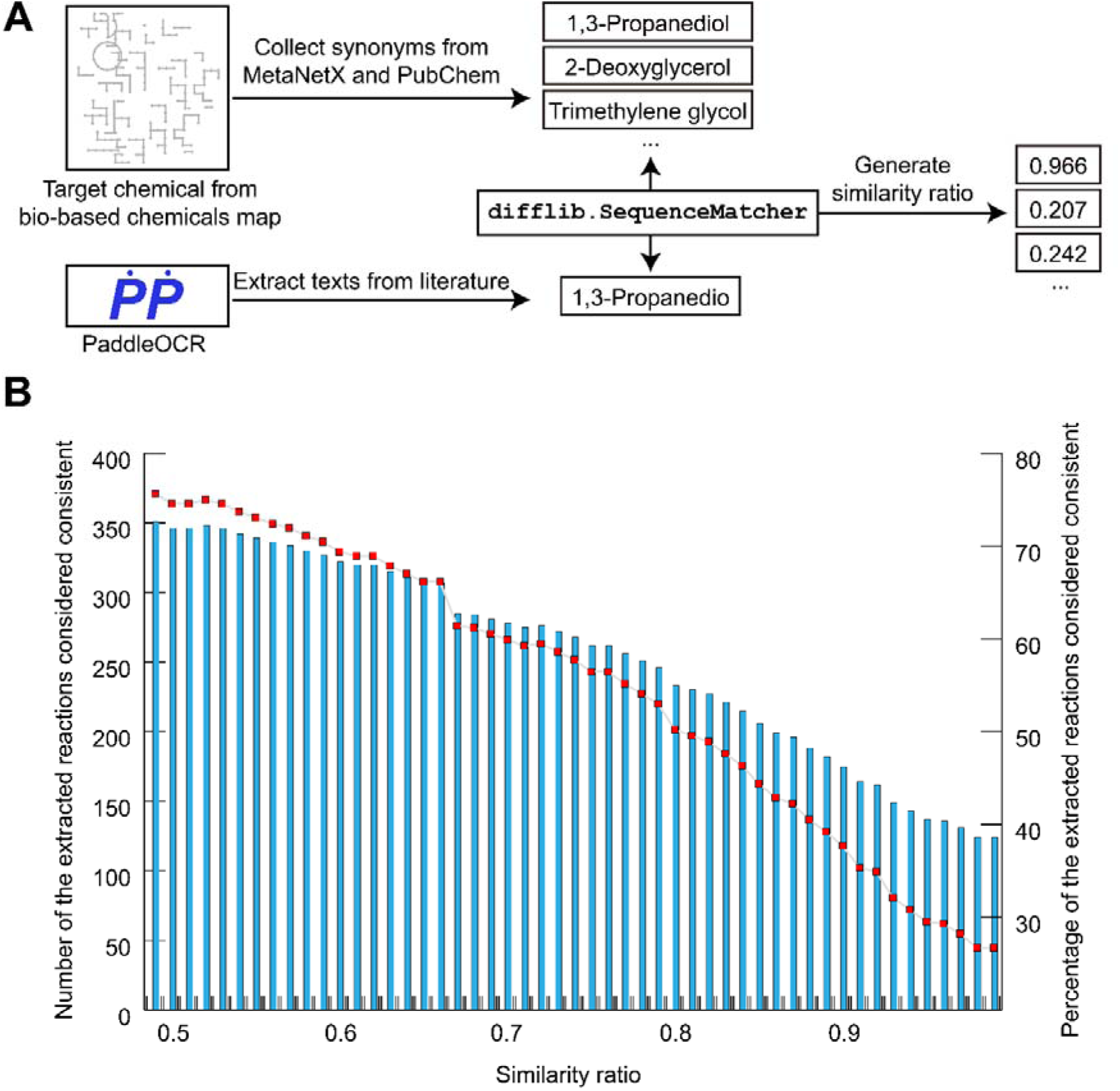
Extraction of biochemical reaction information for target chemicals from the bio-based chemicals map (Jang et al., 2023) by using EBPI. (A) Process of extracting biochemical reaction information using EBPI and evaluating it against the bio-based chemicals map. Texts from EBPI and the bio-based chemicals map were compared by using the `difflib.SequenceMatcher` function in Python. The similarity ratio, ranging from 0 to 1, is an output value from `difflib.SequenceMatcher`, and it was used to determine the consistency level of the EBPI output (i.e., metabolite name). (B) Number and percentage of the extracted reactions considered consistent across varying similarity ratios. The entire target chemicals considered in this analysis were associated with 464 biochemical reactions. Blue bars (on the left y-axis) indicate the number of reactions successfully extracted by EBPI among the 464 biochemical reactions. Red dots (on the right y-axis) indicate the percentage of reactions successfully extracted from the total of 464 biochemical reactions.

For this comparison, EfficientNet (Tan and Le, 2020), a lightweight model for accurate classification of images, was deployed to efficiently select relevant images for the EBPI input. This step was necessary because papers often have irrelevant images (e.g., images not having arrows as well as “non-pathway” images despite having arrows), and such images were discarded by using EfficientNet (Fig. 3A). However, EfficientNet needs to be trained. For this, we retrieved a total of 424 papers using the search term “*E. coli* metabolic pathway” from PMC. Images were retrieved from the collected papers by using a Python package PyMuPDF (https://pypi.org/project/PyMuPDF/) that extracts images from PDF files. Next, the trained Faster R-CNN for EBPI, elaborated above, was used to assign the images as positive and negative data; the positive data are the images with five or more detected arrows, while the negative data are the ones with fewer than five detected arrows. A threshold of five arrows was determined based on the average and median number of arrows in MetaCyc pathways, which were 5.76 and 4.00, respectively (Fig. 3B). The negative data underwent further manual curation to select the additional qualifying images, and completely discard the remaining negative images, such as tables or graphs with arrows (Fig. 3C). As a result, the numbers of positive (“pathway”) and negative (“non-pathway”) images were 273 and 2,069, respectively; both data were used to train EfficientNet.

Next, a maximum of 2,000 papers were collected for each of the 466 chemicals from the bio-based chemicals map (Fig. 3D). Threshold for the maximum number of papers for a target chemical was set to 2,000. If the number of papers searched for a target chemical was below 2,000, all the papers were considered for the EBPI implementation. Here, a total of 74,853 papers were collected for 466 target chemicals. Next, images were retrieved from the collected papers using PyMuPDF if the separate image files were not available for the PMC papers in .gzip format (Methods). The collected images were subsequently filtered by using the trained EfficientNet (Fig. 3E). As a result, 49,846 images were finally selected from an initial set of 677,618 images for 421 chemicals; no images were selected for 45 chemicals because papers were not searched for 39 of these chemicals, and no images passed the selection process using EfficientNet for the remaining 6 chemicals.

Final step is to compare the biochemical reaction information extracted by EBPI (https://zenodo.org/records/11075692) with that available in the bio-based chemicals map. For this comparison, synonyms of the 421 target chemicals were collected from MetaNetX (Moretti et al., 2021) and PubChem (Kim et al., 2023a) because images from the selected papers and the bio-based chemicals map may use different names for the same molecule. Next, texts (i.e., names of substrates and products) from EBPI and the bio-based chemicals map were compared by using the `difflib.SequenceMatcher` function in Python (Fig. 4A); this function compares pairs of hashable sequences, and generates the so-called similarity ratio that ranges between 0 and 1. As a result, the percentage of extracted biochemical reactions considered consistent changes, depending on the a priori set similarity ratio (Fig. 4B). In this study, the similarity ratio was increased from 0.49 to 0.99 with an interval of 0.01. As a result, the percentage of the extracted reactions, which were considered consistent, dropped from 75.6% to 26.7%, and it was 50.2% when the similarity ratio was 0.8. Implementation of EBPI for target chemicals from the bio-based chemicals map revealed factors that affected its performance. First, no papers were retrieved from PMC for 39 target chemicals. For example, a search term “β-Hydroxy-γ-butyrolactone AND metabolic engineering” in PMC did not retrieve any papers for β-hydroxy-γ-butyrolactone, a target chemical covered by the bio-based chemicals map. Lack of such papers negatively affected the performance of EBPI. Second, not all the intermediate metabolites are presented in the bio-based chemicals map due to the limited space. This obviously disables EBPI to extract relevant biochemical reaction information. This can be an inherent limitation of EBPI when extracting pathway information from image data in papers.

### 2.5. Extraction of biochemical reaction information not covered by biological pathway databases

A benefit of using EBPI is the ability to extract biochemical reaction information not only reported in the literature, but also those not covered by biological pathway databases (Fig. 5). To confirm whether EBPI extracts biochemical reaction information currently not covered by biological pathway databases, we selected target chemicals from the bio-based chemicals map that satisfied at least two of the following three criteria: 1) absence of MetaNetX ID for biochemical reactions of a target chemical; 2) absence of a KEGG compound ID for a target chemical; and 3) a similarity ratio below 0.8 between the names of all intermediate metabolites extracted using EBPI and those listed in the bio-based chemicals map for a target chemical. Criteria 1 and 2 suggest a high likelihood that the reactions associated with production and/or consumption of a target chemical are absent in the representative biological pathway databases such as MetaNetX and KEGG. Criterion 3 is to exclude abbreviated reactions that could be incorrectly considered as a novel reaction in the literature figure; for example, glycolysis is sometimes briefly presented as “D-glucose → 2 pyruvate” in a figure, which is not obviously a novel reaction. If all the intermediate metabolite names extracted by EBPI have the similarity ratio below 0.8 in comparison with a list of manually prepared intermediate names, there is a high chance of finding biochemical reactions not covered by the databases. It should be noted that intermediate metabolites associated with glycolysis, pentose phosphate pathway, and TCA cycle were considered for the criterion 3, and their synonyms were also considered when calculating the similarity ratio. As a result of applying the three criteria to the 466 chemicals, 43, 46, and 93 target chemicals were identified to meet the criteria 1, 2, and 3, respectively. Among these, reactions not covered by the databases were found for 1,4-butanediol (meeting the criteria 2 and 3), 2-methylbutyric acid (meeting the criteria 1 and 2), hydroxytyrosol (meeting the criteria 2 and 3), levulinic acid (meeting the criteria 1 and 2), and valerolactam (meeting the criteria 2 and 3) (Fig. 5).

**Fig. 5.**
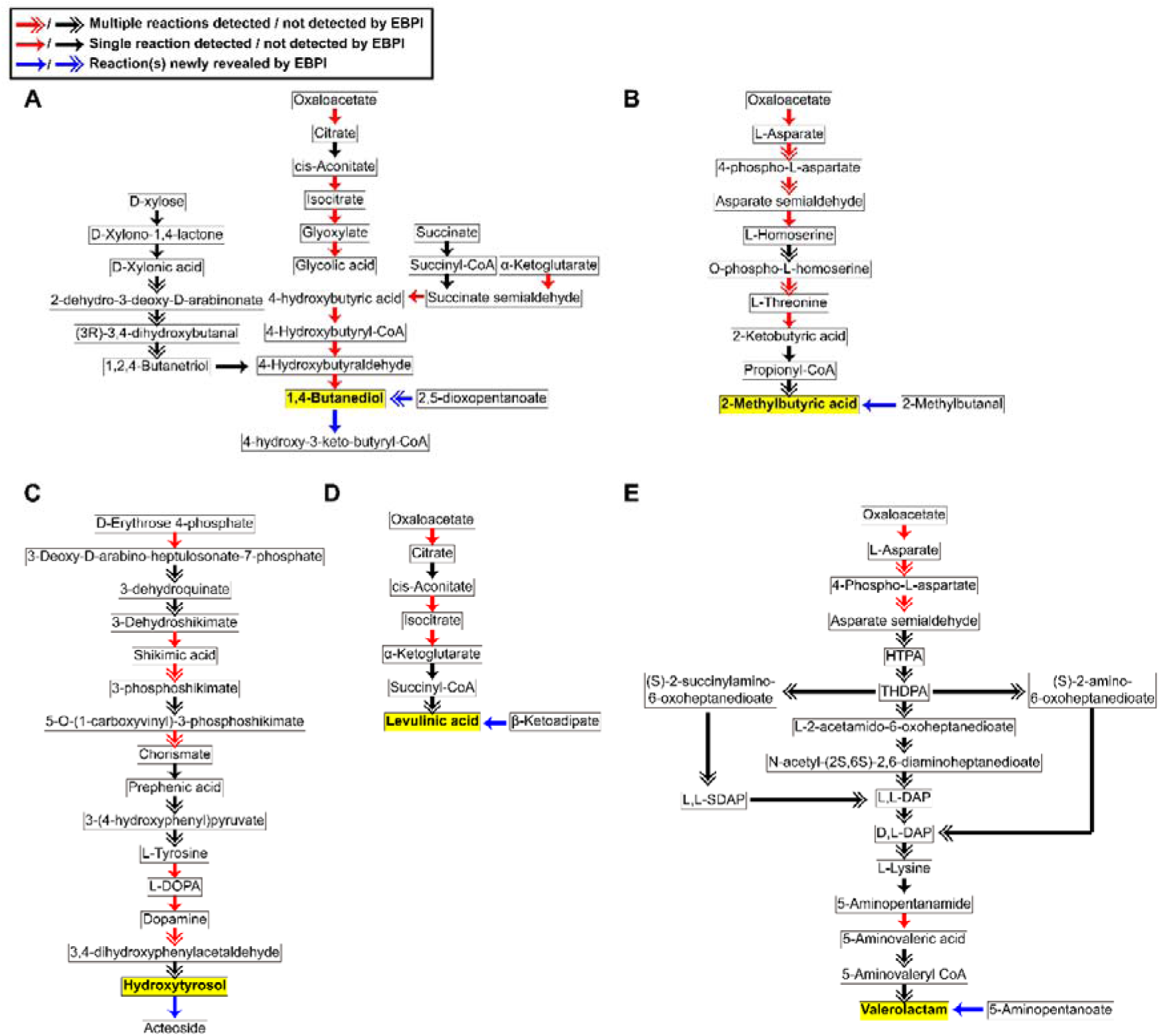
EBPI results uncovering target chemical-specific reactions not cataloged in MetaNetX and/or KEGG databases. Yellow boxes indicate target chemicals having such reactions not covered by KEGG and/or MNXref. The arrow types represent the detection status of reactions by EBPI (see the label in the upper left corner). The following six biochemical reactions, marked with blue arrows, were uncovered from papers on the five different target chemicals. (A) 2,5-Dioxopentanoate to 1,4-butanediol (Shepelin et al., 2018), and 1,4-butanediol to 4-hydroxy-3-keto-butyryl-CoA (Qian et al., 2023). (B) 2-Methylbutanal to 2-methylbutyric acid (Wu et al., 2022). (C) Hydroxytyrosol to acteoside (Li et al., 2022). (D) β-Ketoadipate to levulinic acid (Kim et al., 2023b). (E) 5-Aminopentanoate to valerolactam (Hendry et al., 2021). Abbreviations: D,L-DAP, D,L-diaminopimelate; HTPA, (2S,4S)-4-Hydroxy-2,3,4,5-tetrahydrodipicolinate; L-DOPA, 3,4-Dihydroxy-L-phenylalanine; L,L-DAP, L,L-diaminopimelate; L,L-SDAP, N-succinyl-L,L-2,6-diaminopimelate; and THDPA, 2,3,4,5-tetrahydrodipicolinic acid.

1,4-Butanediol is a widely used chemical in various applications, including fibers, adhesives, textiles, plastic processing, and drugs (Fig. 5A). Due to its extensive applications, it has been an important target for overproduction through metabolic engineering (Kumar et al., 2023). For example, 1,4-butanediol was overproduced by engineering nonphosphorylative pathway in *Escherichia coli* (Shepelin et al., 2018). In this study, two biochemical reactions were newly extracted from the literature: 2-ketoacid decarboxylase (KDC) and alcohol dehydrogenase (ADH) converting 2,5-dioxopentanoate ultimately to 1,4-butanediol (Shepelin et al., 2018); and alcohol dehydrogenase (*calA*, PP_2049, PP_2827), aldehyde dehydrogenase (*peaE*) and oxidoreductase (PP_3208, PP_2737, *fadBA*) converting 1,4-butanediol to 4-hydroxy-3-keto-butyryl-CoA (Qian et al., 2023). Systematic identification of reaction information, which is missing in such representative databases, could help identify additional engineering targets for further optimization of production performance.

The other four chemicals also have industrial values, and relevant biochemical reactions were revealed by using EBPI, which are also not available in MetaNetX and KEGG. 2-Methylbutyric acid and its derivatives (e.g., 2-methylbutanoic acid methyl ester) are primarily used as flavors and food additives due to their distinct smells (Kwon et al., 2000) (Fig. 5B). In this occasion, a biochemical reaction (dihydrolipoyllysine-residue transferase) converting 2-methylbutanal into 2-methylbutyric acid was identified (Wu et al., 2022). Hydroxytyrosol, which is found in olive leaves and olive oil primarily in the form of oleuropein, is utilized for its anti-inflammatory, antioxidant, anticancer, antimicrobial, and antidiabetic properties (Arangia et al., 2023) (Fig. 5C). A biochemical reaction (UDP-glucose glucosyltransferase) was extracted that was reported to convert hydroxytyrosol to acteoside (Li et al., 2022). Levulinic acid serves as a versatile building block chemical with various applications, including pharmaceuticals, food, cosmetics, rubber, and plastics (Kumar et al., 2020) (Fig. 5D). A biochemical reaction (acetoacetate decarboxylase) converting β-ketoadipate to levulinic acid was discovered by using EBPI (Kim et al., 2023b). Lastly, valerolactam serves as an important precursor chemical for synthesizing bioplastics and biopolyamides such as clothes, architecture, and disposable goods (Cheng et al., 2021) (Fig. 5E). A reaction catalyzed by lactam synthase converting 5-aminopentanoate into valerolactam was identified (Hendry et al., 2021). These are several examples demonstrating the benefits of using EBPI for extracting reactions associated with target chemicals that are not covered by the presentative biological pathway databases.

## 3. Discussion

In this study, a machine learning framework EBPI was developed that extracts information on biochemical reactions from images in the biological literature (Figs. 1 and 2). EBPI processes images showing biological pathways, including metabolites, genes and proteins, to generate the pathway information in an editable, tabular format. For this, Faster R-CNN and PaddleOCR were implemented to detect arrows and texts within pathway images, respectively (Fig. 2). Upon development, the performance of EBPI was tested by comparing the extracted biochemical reaction information for 466 target chemicals with those from the bio-based chemicals map (Figs. 3 and 4). Also, EBPI was used to uncover reactions associated with any of the 466 target chemicals, which were not documented in MetaNetX and KEGG, two representative biological pathway databases (Fig. 5). These simulation studies using EBPI demonstrated its potential in identifying biological pathways for a wide range of industrially valuable chemicals, including those not covered by the representative biological pathway databases. In particular, proper use of EBPI in the early phase of a metabolic engineering project can be beneficial, especially for target chemical selection and design of biosynthetic pathways. For example, the EBPI-generated output helps in estimating the difficulty level of a metabolic engineering project when considering target chemical candidates; a candidate project is likely to be more feasible if a greater number of biochemical reactions are extracted for a specific target chemical. The EBPI output can also help narrow down the target reactions generated by retrobiosynthesis; a challenge of retrobiosynthesis is the generation of a large number of candidate biosynthetic pathways (Kim et al., 2021).

While developing EBPI, several challenges have become clear that need to be addressed for its refinement and application. A primary challenge is the validation of pathway information extracted from the literature, distinguishing between experimentally validated and hypothetical ones. This validation would require manual inspection of the EBPI’s output. Implementing state-of-the-art language models can be one option to streamline this time-consuming process. Next, continued improvement in the performance of object detection models (e.g., Faster R-CNN), OCR models (e.g., PaddleOCR), and language models (e.g., BioBERT used in this study) will further enhance the extraction performance of EBPI. Faster R-CNN, PaddleOCR and BioBERT were chosen for this study due to their high performance and accessibility within research communities. However, future versions or more sophisticated models, including commercial ones, may be considered if the extraction performance of EBPI needs to significantly exceed what is presented here. Successfully addressing these challenges will enhance EBPI’s functionality for a wider range of biotechnological applications.

## 4. Material and methods

### 4.1. Preparation of pathway images for arrow detection

Finite Automata Domain, Online Handwritten Flowchart Dataset, and FlowChart Domain offered data labels for the bounding boxes, but such labels were not available for FR-DETR, MetaCyc, and the test datasets (i.e., 100 pathway images) retrieved from the 89 papers (Fig. 1A,B). Therefore, bounding box were manually labeled using labelImg 1.8.1 (https://github.com/HumanSignal/labelImg), which generates coordinates for a selected bounding box in an image. Rotation and random crop of the original images were conducted using Albumentations 1.3.0 (https://www.mdpi.com/2078-2489/11/2/125) to generate more diverse arrows with different directions and sizes.

### 4.2. Preparation of pathway images for text classification

For the text classification, 15,101 genes, 21,417 proteins including polypeptides and protein complexes, and 18,972 compounds were collected from MetaCyc (Fig. 1C). The data were collected from MetaCyc (August, 2023) through “List of All Genes”, “List of All Polypeptides”, “List of All Protein Complexes” and “List of All Compounds” in “Search Genes, Proteins or RNAs” and “Search Compounds”. These terms were used to classify the texts detected from images to “gene”, “protein” or “others”.

### 4.3. Optimization of object detection models for arrows

Prediction performances of Faster R-CNN (Ren et al., 2016) (both V1 and V2) and DETR (Carion et al., 2020) (both original and deformable versions) were examined using the test dataset using the 100 pathway images (Fig. 1B and Table 1). These object detection models were subjected to hyperparameter optimization. For this, following values were considered for each of the main hyperparameters: [8, 16] for `batch_size`; [10^-2^, 10^-3^, 10^-4^, 10^-5^] for `lr`; [1, 5] for `max_norm`; and [10^-3^, 10^-4^] for `weight_decay`. Threshold for the confidence score of Faster R-CNN and DETR was set to 0.5 when determining whether to select a detected arrow. Selected arrows were considered to calculate the mean average precision (mAP) score. APs were calculated using intersection over union (IoU) ranging from 0.50 to 0.95 with an increment of 0.05. Finally, all the calculated APs were averaged to compare the overall performance of Faster R-CNN (both V1 and V2) and DETR (both original and deformable versions).

Hyperparameters of the best-performing Faster R-CNN V1 (`fasterrcnn_resnet50_fpn_v1`) were 10^-5^, 10^-3^, 1 and 8 for `lr`, `decay`, `clip_grad_norm` and `batch_size`, respectively. For the optimal Faster R-CNN V2 (`fasterrcnn_resnet50_fpn_v2`), its hyperparameters were 10^-4^, 10^-4^, 5 and 8 for `lr`, `decay`, `clip_grad_norm` and `batch_size`, respectively. Similarly, for DETR, the best set was 10^-5^, 10^-4^, 5 and 8 for `lr`, `decay`, `clip_grad_norm` and `batch_size`, respectively. Finally, for Deformable DETR, the best set was 10^-5^, 10^-4^, 5 and 4 for `lr`, `decay`, `clip_grad_norm` and `batch_size`, respectively. Among these configurations, `fasterrcnn_resnet50_fpn_v2` showed the best performance, achieving mAP of 0.433 at a confidence score of 0.5 (Table 1).

### 4.4. Detection of arrow directions within a pathway image

After bounding boxes of arrows were detected using an object detection model, OpenCV 4.7.0 (https://github.com/opencv/opencv) was used to identify connected objects within these bounding boxes (Fig. 2A). The aforementioned function helps determine the longest connected objects within a bounding box, enabling the removal of irrelevant objects (e.g., any images that are not part of an arrow). Subsequently, OpenCV was used to skeletonize the arrow and extract its end-points. Finally, the end-points were grouped using *k*-means clustering to detect the head and tail of the arrow. This procedure led to the identification of the direction and reversibility of a detected arrow.

### 4.5. Optimization of BioBERT for text classification

BioBERT 1.2.0 (Lee et al., 2020) was used in this study to classify the detected texts into “gene”, “protein” or “other” (Fig. 2B). BioBERT was trained using the collected MetaCyc terms and the papers searched through PMC (Fig. 1C). Following values were explored using grid search: [4, 8, 16, 32] for `batch_size`; and [10^-4^, 10^-5^, 10^-6^] for `lr`. As a result, `batch_size` and `lr` were set to 16 and 10^-5^, respectively, resulting in an accuracy of 0.975 for the test dataset.

### 4.6. Detection of texts within a pathway image

The pre-trained PaddleOCR 2.6.0 (https://github.com/PaddlePaddle/PaddleOCR) was used in this study to detect texts in an image (Fig. 2B). PaddleOCR was used in this study because it appeared to outperform EasyOCR (https://github.com/JaidedAI/EasyOCR), another open-source OCR model, for the text detection. Texts detected using PaddleOCR were subsequently classified into “gene”, “protein” or “others” by using the trained BioBERT. The relative locations of the classified texts were calculated with respect to the head and tail of an arrow, which then allowed for the generation of biochemical reaction information.

### 4.7. Preparation of pathway images for 466 target chemicals in the bio-based chemicals map

To evaluate the performance of EBPI, pathway information in the bio-based chemical map (Jang et al., 2023) was used, which covers a total of 466 chemicals (Figs. 3 and 4). Among the 466 chemicals, relevant papers could be searched for 455 chemicals. BioPython (https://biopython.org/) and FTP were used to download the papers for these 455 chemicals. BioPython was used to extract PMCID of up to 2,000 articles for each of the 455 chemicals; a target chemical name and a word “metabolic engineering” were used as queries in BioPython. Subsequently, PMCID was used to find corresponding papers from “comm_use_file_list.csv” and “non_comm_use_pdf.csv” in the NCBI FTP site (https://ftp.ncbi.nlm.nih.gov/pub/pmc/). If the matching PMCID was available in “comm_use_file_list.csv”, relevant image files in tar.gz compressed format could be downloaded from the FTP server. For “non_comm_use_pdf.csv”, entire articles were downloaded in PDF format if the matching PMCID was available. In this case, an additional process was necessary to prepare pathway images from the PDF files. For this, PyMuPDF 1.22.5 (https://pypi.org/project/PyMuPDF/1.22.5/) was used to extract the pathway images from the PDF files. Subsequently, EfficientNet was trained using 2,342 images from the 424 papers collected from PMC, which were searched using the term “*E. coli* metabolic pathway”. Finally, a trained EfficientNet model, `EfficientNet-b0`, was used to select the images that fit the objective of this study.

To retrieve information presented in the bio-based chemicals map for each of the target chemicals, MetaNetX (Moretti et al., 2021) IDs of metabolites were downloaded from http://systemsbiotech.co.kr/. Each MetaMetX ID of metabolites was converted to the ChEBI (Hastings et al., 2016) ID, and “description” information in “chem_xref.csv” from the MNXref namespace (Moretti et al., 2021) (https://www.metanetx.org/mnxdoc/mnxref.html; version 4.4). These ChEBI IDs and “description” information were used to collect metabolite synonyms from ChEBI and PubChem (Kim et al., 2016) (October, 2023). Synonyms were considered when comparing the data from the bio-based chemicals map with the pathway information extracted using EBPI.

## CRediT authorship contribution statement

**Mun Su Kwon:** Conceptualization, Data curation, Formal analysis, Investigation, Methodology, Software, Validation, Visualization, Writing-original draft, Writing-reviewing & editing. **Junkyu Lee:** Conceptualization, Data curation, Formal analysis, Investigation, Methodology, Software, Validation, Visualization, Writing-original draft, Writing-reviewing & editing. **Hyun Uk Kim:** Conceptualization, Data curation, Funding acquisition, Project administration, Resources, Software, Supervision, Writing-review & editing.

## Acknowledgments

This study was supported by the ERC Center of the National Research Foundation (NRF) funded by Ministry of Science and ICT (NRF-2022R1A5A1033719). This study was also carried out with the support of “Cooperative Research Program for Agriculture Science and Technology Development (Project No. PJ01577901)” from Rural Development Administration, Republic of Korea.

## Data availability

The source codes of EBPI are available at https://github.com/kaist-sbml/EBPI. Training and validation datasets for training Faster R-CNN (Fig. 1A), a test dataset for the trained model (Fig. 1B), and EBPI results for the 466 target chemicals from the bio-based chemicals map (Fig. 4) are available at https://zenodo.org/records/11075692.

